# Intrinsic ion dynamics underlies the temporal nature of resting-state functional connectivity

**DOI:** 10.1101/2025.11.08.687387

**Authors:** Oscar C. González, Pavel Sanda, Jaroslav Hlinka, Maxim Bazhenov, Giri Krishnan

**Author notes:** O.C.G. and P.S. contributed equally to this work. M.B. and G.K. are co-senior authors.

## Abstract

The neural mechanisms underlying the emergence of functional connectivity in resting-state fMRI remain poorly understood. Recent studies suggest that resting-state activity consists of brief periods of strong co-fluctuations among brain regions, which reflect overall functional connectivity. Others report a continuum in co-fluctuations over time, with varying degree of correlation to functional connectivity. These findings raise the critical question: what neural processes underlie the temporal structure of resting-state activity? To address this, we used a biophysically realistic whole-brain computational model in which resting-state activity emerged from temporal variations in the ion concentrations of potassium (K^+^) and sodium (Na^+^), intracellular chloride (Cl^-^), and the activity of the Na^+^/K^+^ ATPase. The model reproduced transient periods of high co-fluctuations, and the functional connectivity at different co-fluctuation levels correlated to varying degrees with the connectivity measured over the entire simulation, in line with experimental observations. The periods of high co-fluctuations were aligned with large changes in extracellular ion concentrations. Furthermore, critical parameters governing ion dynamics strongly affected both the timing of these transient events and the spatial structure of the resulting functional connectivity. The balance of excitatory and inhibitory activity further modulated their frequency and amplitude. Together, these results suggest that intrinsic fluctuations in ion dynamics could serve as a plausible neural mechanism for the temporal organization of co-fluctuations and resting-state functional connectivity.

## Introduction

Resting-state fMRI activity is widely used to assess brain function in both healthy and disease conditions (1, 2). In particular, the functional connectivity (FC) derived from blood oxygen level-dependent (BOLD) activity between different brain regions has been used to identify the neural basis of cognitive processes (3, 4) and alterations in diseases (5). Coherent fluctuations across distinct brain areas, typically quantified by Pearson correlation, give rise to functional connectivity, which reflects the interactions between distributed brain regions at rest (6, 7).

Across many studies, a single FC measure is computed for the entire scanning session. However, this approach ignores the temporal variations in FC, even though electrophysiological recordings have shown that neural activity is dynamic and undergoes large fluctuations over time (8-12). More recent work has focused on the temporal variations in FC and has indeed found that FC varies across time (13). Sliding window analysis (14) has revealed transient “spontaneous events” involving multiple subnetworks (15). More recently, single frame-based methods have been applied to examine moment-to-moment variations in resting-state FC (16). In these works, an “edge-centric” analysis, where pairwise regional co-fluctuations (edges) are measured at single time frame resolution, was used (17-19). This approach further demonstrated that FC varies over time, with ongoing resting-state activity punctuated by brief periods of widespread highly correlated activity (16).

Several studies examining FC now suggest that FC varies over time, however, many properties of time-resolved FC remain a contested topic (13). One issue concerns the contribution of brief events to the overall FC. Studies using the single frame analyses have found that brief periods of high co-fluctuation are strongly correlated with the overall FC measured across the entire recording period (19). In contrast, other studies have reported a monotonic relationship between the strength of co-fluctuation and the overall FC (20, 21). Further, statistical models have suggested that static FC alone could drive such brief high co-fluctuation events (22). Collectively, these findings have led to an ongoing debate whether co-fluctuations represent genuine neural events (23-26) or arise from experimental factors (e.g. head motion) or statistical artifacts (e.g. variability in sampling from static FC) (20-22, 27).

A complementary approach to studying the dynamics of FC is to use bottom-up methods based on the modeling of spontaneous neural activity, and infer the derived statistical measures from the simulated data. This approach assumes that neural signals drive BOLD activity, with artifacts such as head motion or noise added on top. The level of detail captured by these methods ranges from coarse-grained field or mass models that describe regional activity to single neuron level models using spiking networks (28-30), with each scale contributing distinct mechanisms (31).

This new work focuses on a detailed biophysical model that generates infra-slow resting-state activity due to ion dynamics (32). Our research was motivated by the following two questions: (a) Does the detailed model reproduce the fast time-scale co-fluctuation dynamics observed in experiments? (33) and (b) What neural mechanisms determine the temporal variations in co-fluctuation and fMRI activity.

Using a large-scale computational network model based on the human connectome, we show that co-fluctuations in resting-state fMRI activity, consistent with observations from experiments, are linked to fluctuations in extracellular K^+^ concentrations ([K^+^]_o_). To establish causality, we demonstrate that altering [K^+^]_o_ ion dynamics or the Na^+^ /K^+^ pump alters the spatial extent and distribution of co-fluctuations. Furthermore, we show that the balance of excitatory and inhibitory network connectivity modulates the co-fluctuations. Overall, our computational model offers a testable hypothesis for the neural mechanism underlying co-fluctuation events, providing a framework for future experimental validation.

## Results

In this study, we used a biophysically realistic cortical network model based on our previous work (34) to investigate the neural mechanisms underlying resting-state infra-slow fluctuations. The model (see Methods for details) consisted of detailed two-compartment excitatory pyramidal (PY) neurons and inhibitory interneurons (IN) with Hodgkin-Huxley kinetics. Neurons were synaptically connected through AMPA, NMDA, and GABA-A synapses. Each neuron received a random Poisson drive to capture stochastic afferent input. The model incorporated dynamically varying ion concentrations for the major ion species, including K^+^, Na^+^, Cl^-^, and Ca^2+^, to mimic *in vivo*-like ionic dynamics.

Figure 1A illustrates the modeled ion concentrations, membrane ion channels, ion exchangers/pumps, and extracellular astrocytic K^+^ buffering. A “cluster” or “node” of neurons was used to model an individual brain region, and consisted of a population of 50 excitatory (PY) and 10 inhibitory (IN) neurons. Neurons within a cluster were connected through synapses and coupled by ion diffusion within the shared extracellular space. An example schematic of a single cluster is shown in Figure 1B. Multiple clusters were modeled and connected via long-range feedforward excitatory synaptic connections (E-to-E and E-to-I) (Figure 1C) to form a network of connected brain regions. The long-range network connectivity was based on either CoCoMac brain data (35) for macaque brain simulations or DW-MRI structural connectivity matrices (36) for human brain simulations. Adjacency matrices for human and macaque simulations are shown in Figure 1D and 1E where color indicates the relative strength of the feedforward excitatory connections.

**Figure 1.**
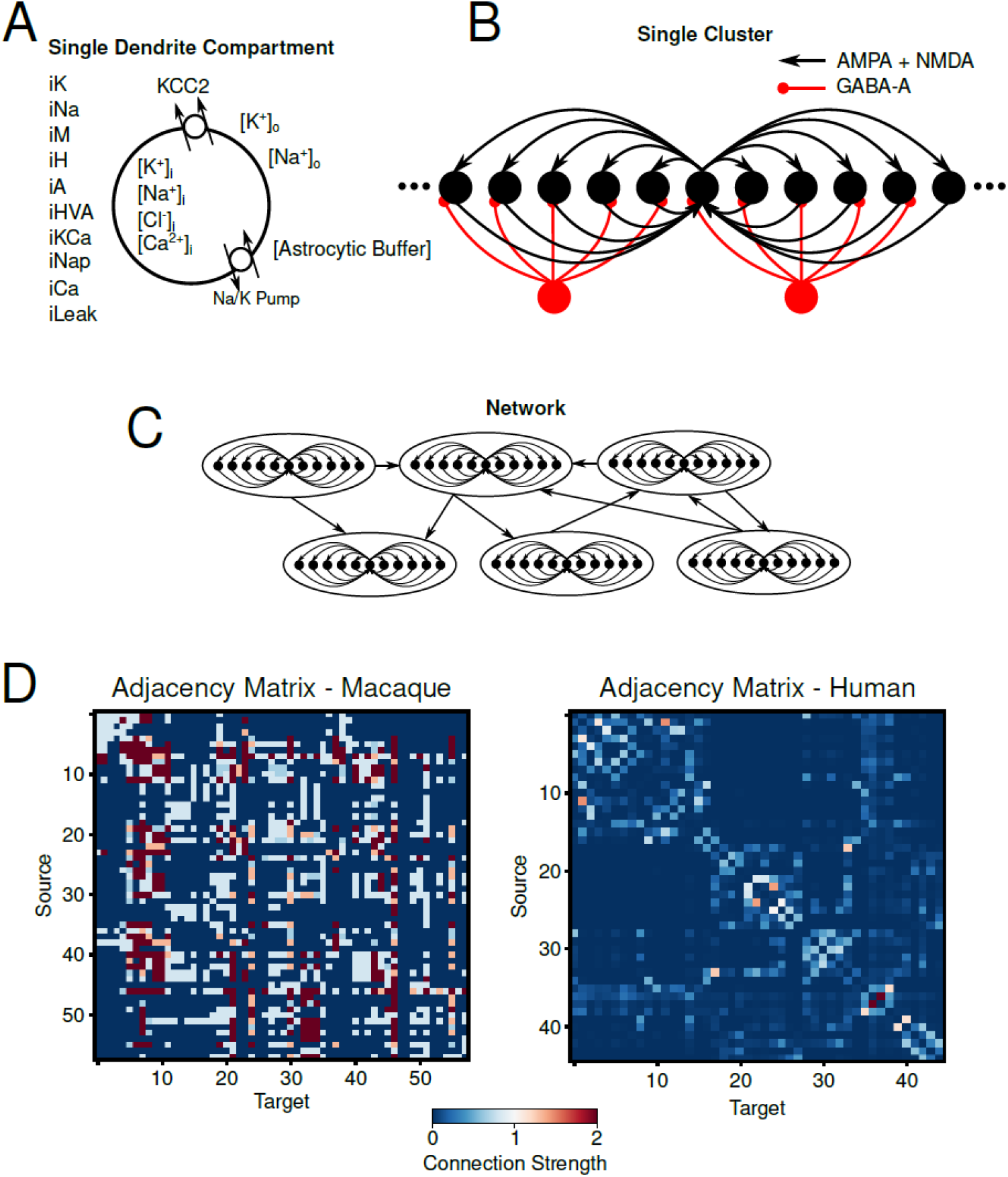
Basic model schematic from single compartment to global connectivity. A, Schematic of single dendritic compartment including a list of intrinsic ionic currents included in the compartment (left), ionic species modeled with dynamics concentrations, KCC2 cotransporter and Na^+^/K^+^ pump, and astrocytic buffer of [K^+^]_o_. B, Schematic of a single cluster of excitatory and inhibitory neurons used to model individual brain regions in the whole network. Black/red circles represent excitatory/inhibitory neurons. Excitatory synapses are mediated by AMPA/NMDA receptors while inhibitory synapses are mediated by GABA-A. C, Schematic representation of how individual clusters for the whole network model. Clusters are connected via feedforward excitation. D, Adjacency matrices for Macaque (left) and Human (right) network simulations. Color indicates the relative strength of the feedforward connections between clusters.

In a network comprising of 58 brain regions of the macaque brain interconnected via feedforward excitation, the average network firing rate exhibited spontaneous infra-slow fluctuations (Figure 2A). As previously reported (32), these fluctuations were also present in the network-wide mean [K^+^]_o_ concentration, intracellular sodium concentration ([Na^+^]_i_), and Na^+^/K^+^ pump current (Figure 2A). Figure 2B shows the corresponding power spectra for each of these signals, displaying a prominent ∼0.025 Hz peak, characteristic of resting-state infra-slow fluctuations commonly observed in brain networks (37, 38). Similar dynamics were observed in a network consisting of 45 brain regions of the human brain simulated using DW-MRI derived connectivity (Figure 2D/E).

**Figure 2.**
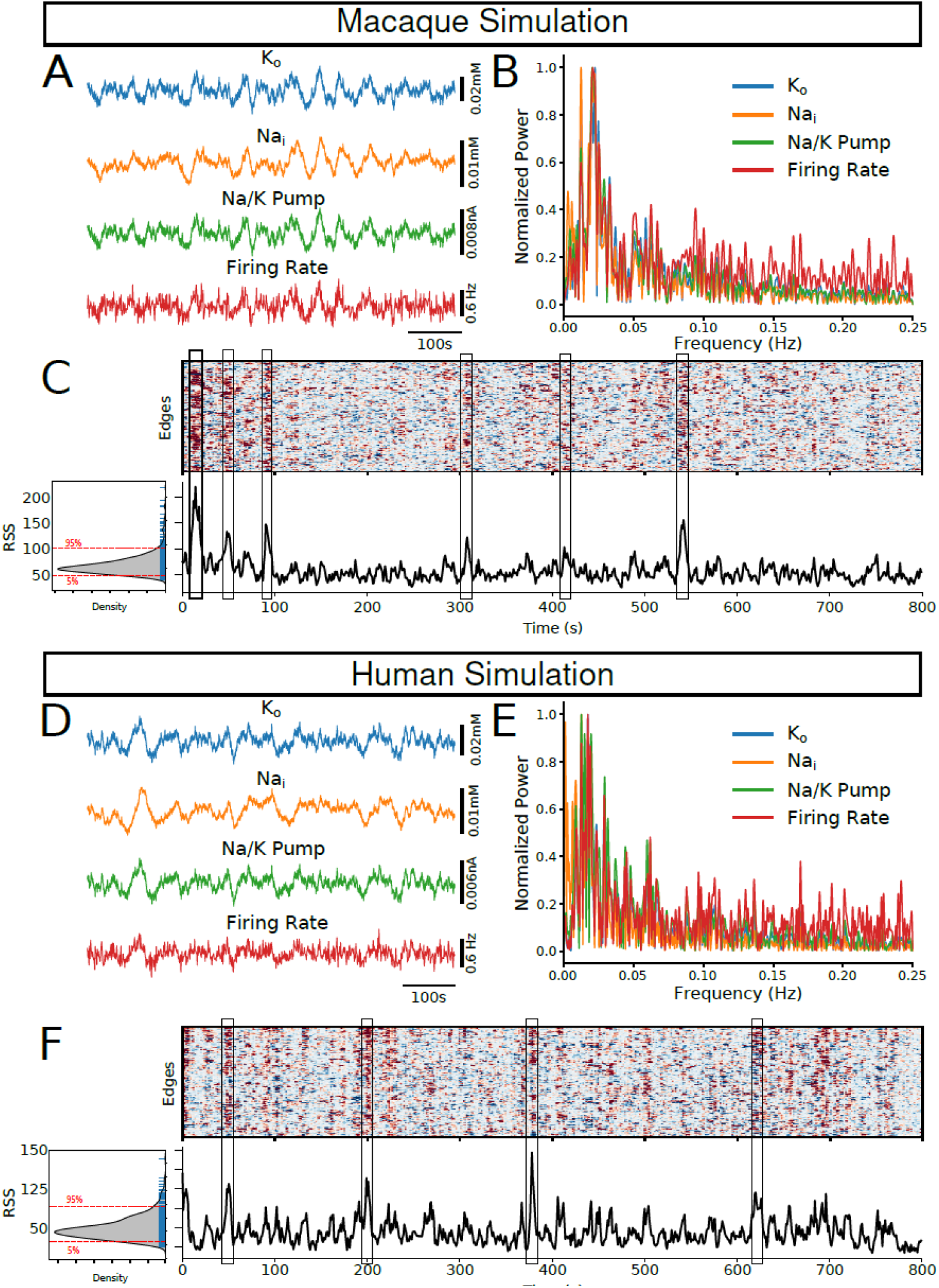
Model with dynamic ion concentrations captures peaks in moment-to-moment co-fluctuations. A/D, Network average [K^+^]_o_, [Na^+^]_i_, Na^+^/K^+^ pump, and firing rate showing characteristic infra-slow fluctuations in both Macaque (A) and Human (D) simulations. B/E, power spectra for the time series in A/D showing peaks around the infra-slow frequencies (0.01-0.05Hz). C/F, Top heatmap shows the computed Edge-based Time Series (EST) for both Macaque and Human network simulations. Bottom, the corresponding Root-Sum Squared (RSS) computed from the EST reveals brief periods of high co-fluctuations throughout the networks. Left, inset shows the distribution of RSS values showing a right skewed distribution.

### The temporal evolution of resting-state activity in the model aligns with experimental observations

To examine temporal variations in functional connectivity during resting periods, we employed a moment-to-moment estimation method previously used in experimental works (20, 39). In this approach, moment-to-moment co-fluctuations are quantified using edge-based time series (ETS), computed for each edge (pair of regions) as the product of z-scored time series from the two regions. Since the Na^+^/K^+^ pump is responsible for the majority of the oxygen consumption in the brain, we used Na^+^/K^+^ pump current as a proxy for the BOLD signal for each region to compute the ETS in this work. Intuitively, ETS captures the relative synchronization between regions rather than differences in absolute signal intensity. Figure 2C,F shows the resulting ETS (top) and the corresponding root-sum squared (RSS) signal of the ETS (bottom) for both macaque and human simulations. Consistent with (19), we found brief periods of high amplitude co-fluctuations, as demonstrated by peaks in the RSS signal (Figure 2C,F, bottom). Additionally, the distribution of RSS values resembled the skewed distribution for RSS signals reported for human fMRI BOLD recordings (Figure 2C,F, bottom).

Experimental studies have shown that periods of high amplitude co-fluctuations exhibit strong correlations with time-averaged rsFC (17-19). In other words, short bursts of high co-fluctuation between regions – moments when many areas are synchronously active – reproduce much of the structure of the full rsFC. Thus, the brain’s overall connectivity pattern appears to be driven by brief, high-amplitude co-fluctuation events. To test whether our model captured this phenomenon, we computed the time-average rsFC of macaque and human simulations, and compared them to the rsFC estimated only from the top 5% of time points in the RSS signal (95th percentile). In other words, the time-averaged rsFC was computed using the entire time series, whereas the “ETS-based” FC was derived from the segments of the time series showing the highest synchrony across regions (top 5% of the RSS signal). Figure 3A/B shows the macaque/human time-averaged rsFC (left) and the mean ETS-based FC from the top 5% RSS time points (middle). The time-averaged rsFC and the ETS-based FC from only the top 5% RSS periods had high similarity, as previously reported in humans. We found that the strongest correlations between time-averaged rsFC and ETS-based FC occurred at the highest RSS amplitude time points (Figure 3A/B, right).

**Figure 3.**
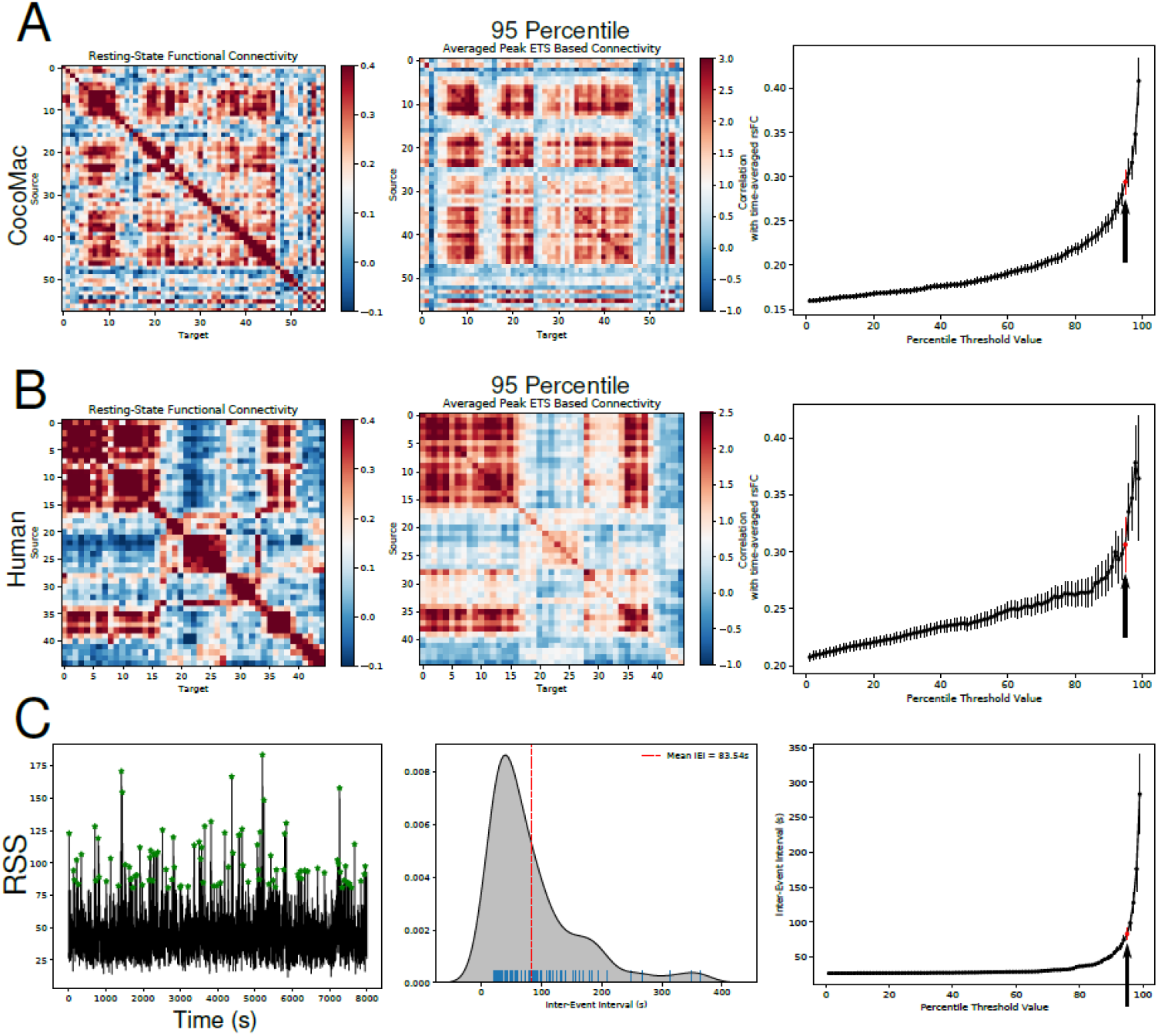
Highest amplitude peaks in RSS correlate strongly with time-averaged rsFC. A/B, Left shows the time-averaged resting-state functional connectivity (rsFC) computed across entire simulated time for Macaque (A) and Human (B) networks. Middle panels show the average ETS-based FC at peaks in RSS time series detected using a 95-percentile threshold. Right panels show the correlation between the rsFC (left panel) and the ETS-based FC (middle panel) as a function of percentile thresholds used to detect RSS peaks. Arrow indicates the 95-percentile threshold condition. C, Left panel shows the RSS time series with detected peaks using a 95-percentile threshold (green stars). Middle panel shows the distribution of inter-event intervals of detected RSS peaks. Right panel shows the interevent interval as a function of percentile threshold used to detect RSS peaks. Arrow indicates the 95-percentile condition.

Next, we asked whether the inter-event interval of high amplitude co-fluctuations reflects the infra-slow timescale of the resting-state fluctuations (Figure 3C). We found that the inter-event intervals between high amplitude co-fluctuations (>=top 5%) displayed timescales similar to endogenous resting-state fluctuations (Figure 3C, middle/right). The distribution of the inter-event intervals had a long tail extending up to 400s. This suggests that the generation of high-value RSS events is irregular and may involve spontaneous accumulation of intrinsic and ionic changes driven by stochastic network activity. We further investigate these cellular and ionic mechanisms in subsequent experiments.

### Co-fluctuations of resting-state in the model require dynamically varying [K^+^]_o_

As the results in Figure 3 suggest, in a model with dynamic ion concentrations, moment-to-moment activity co-fluctuations occur spontaneously and contribute to the generation of the time-averaged rsFC, leading us to investigate the role of neuronally-generated infra-slow activity by blocking ion dynamics. Based on our previous work, where we identified [K^+^]_o_ dynamics as a critical component (40), here we primarily investigated the impact of varying [K^+^]_o_ dynamics. We simulated a human network (for an extended period of 2.5 hours of simulated fMRI signal) in 4 different conditions – normal, fixed [K^+^]_o_, removed global connectivity (G) (to disentangle contribution of connectivity), and finally both fixed [K^+^]_o_ and removed G. The effect is visible in Figure 4A/B, where fixing [K^+^]_o_ decreased variability in both firing rate and Na^+^/K^+^ pump current (Figure 4A/B, second panel), but not their relative levels (due to relative network hubness of each node). Blocking G kept variability but removed the relative levels, and put every brain area on the same basic level (Figure 4A/B, second panel). Blocking both combined the effect (fourth panel of Figure 4A/B).

**Figure 4.**
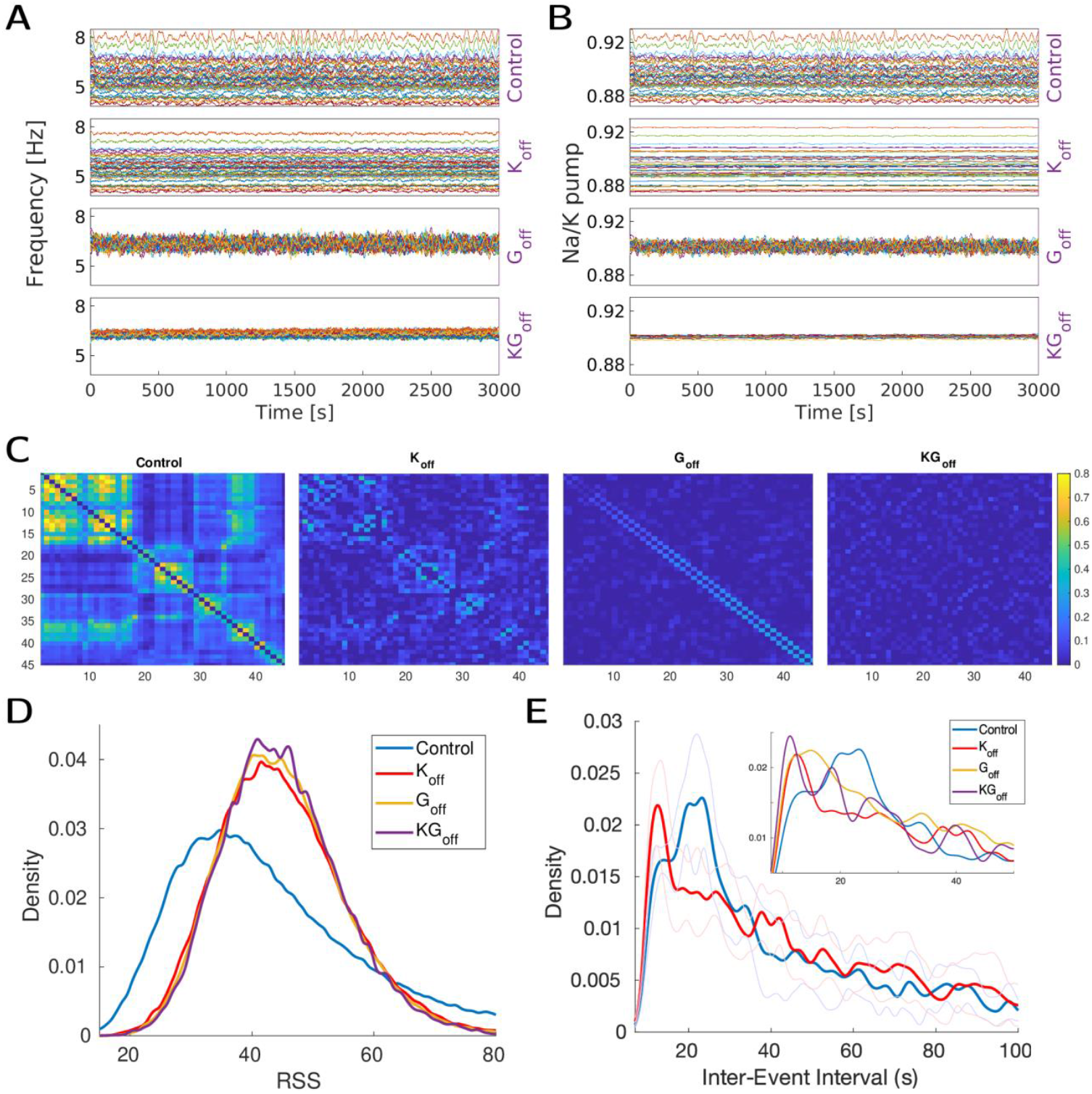
Effect of potassium dynamics on the Inter-event intervals. In all panels 4 conditions are considered: Control: baseline human connectome model. K_off_: model with removed potassium dynamics. G_off_: Model when long-range connections are removed. KG_off_: G_off_ and K_off_ combined. Top. Underlying physiological activity: A, average firing rate of each region (moving average, window 30 [s]). B, average Na^+^/K^+^ pump activity of each region (moving average, window 30 [s]). C, Corresponding functional connectivity for 4 respective conditions. Bottom. RSS-based measures: D, Histogram of raw RSS values for each condition. E, Inter-Event interval length histogram for two major conditions - Control and K_off_ (shaded lines indicate standard deviation, 14 trials). Inset shows all 4 conditions for comparison.

From an fMRI perspective (Figure 4C), blocking G disrupts FC structure (which is partly driven by structural long-range connectivity), while fixing [K^+^]_o_ keeps shallow contours (connectivity for fast interactions is still there, but the major oxygen consumer – Na^+^/K^+^ pump – and driver for slow BOLD signal is severely affected). From the perspective of the RSS measurements (Figure 4D/E), we see changes in the RSS distributions, and a separation of co-fluctuation inter-event intervals between normal and test scenarios, with fixed [K^+^]_o_ resulting in a more pronounced effect. This suggests that the potassium mechanism, being the main driver for the infra-slow activity in the biophysical model (32), impacts co-fluctuation event statistics and influences their occurrence.

### Co-fluctuations are coupled with ion fluctuations

To further understand the temporal relationship between co-fluctuations and ion concentrations/neural activity, we examined periods around co-fluctuation peaks using peak-triggered averaging (Figure S1). We identified two different neural activity patterns that occur during the co-fluctuation events. In the first case (Figure S1B, left), the RSS peak was associated with a significant reduction in firing rate and [K^+^]_o_. The Na^+^/K^+^ pump current had low values during these peaks. In the second case (Figure S1B, right), the RSS peak was associated with increased firing rate and [K^+^]_o_. In both cases, there was a small oscillatory nature to the RSS and ion dynamics, as the peak corresponding to the reduction of firing rate was followed or preceded by a peak with increase in firing rate (similarly for the peak with increased firing rate). This was further verified when we examined the distribution of simulated BOLD, firing rate and [K^+^]_o_ for different RSS values. High RSS values had distributions (Figure S1C) which are significantly higher or lower compared to lower RSS values in all 3 variables. The overall average (Figure S1D) captured only the increase in the firing rate and BOLD, suggesting the number of instances with an increase in firing rate and BOLD was higher than the reduction.

### Accumulation of ionic concentration is required for spontaneous fluctuations

The fluctuations of ion dynamics are both dependent on and influence the neural activity. It undergoes a period of accumulation where it increases and periods of reduction when the accumulated ions are removed through active processes such as Na^+^/K^+^ pump activity. Thus, to investigate further how local ionic fluctuations regulate global dynamics, we altered the strength of the Na^+^/K^+^ pump current locally in each brain region in a model with simulated human connectivity. Reducing the Na^+^/K^+^ pump current will result in lower clearance of the extracellular ions including [K^+^]_o_ ions, while increasing Na^+^/K^+^ pump strength will lead to higher clearance and lower amplitude in ion fluctuations.

Figure 5A shows the average [K^+^]_o_ across all brain regions resulting from different amounts of Na^+^/K^+^ current strength. Reduction of Na^+^/K^+^ pump strength (e.g. 90% pump strength) resulted in large amplitude [K^+^]_o_ fluctuations (Figure 5A). This was due to a slowdown in the ability of the pump to respond to the gradual accumulation of extracellular K^+^ resulting in [K^+^]_o_ reaching higher levels, and thereby increasing the duration of the gradual discharge of accumulated [K^+^]_o_. Alternatively, increasing pump strength (e.g. 120% pump strength) resulted in a much more efficient clearance of [K^+^]_o_ and smaller [K^+^]_o_ fluctuations (Figure 5A). As changes to the rise and decay dynamics of [K^+^]_o_ could impact properties of co-fluctuations in network activity, we examined the effect of Na^+^/K^+^ pump strength on ETS and RSS distributions. ETS computed for low pump strengths (e.g. 90%) showed more pronounced periods of co-fluctuations in network activity (Figure 5B, left). The corresponding RSS time series showed wider RSS peaks than networks with stronger Na^+^/K^+^ pump strengths (e.g. 100% or 110%, Figure 5B middle/right). Similarly, we observed wider RSS distributions for networks with lower pump strengths (Figure 5C, left). We next asked how changes to Na^+^/K^+^ pump strengths could drive the observed differences in RSS and co-fluctuations in network activity. We found that RSS peaks in lower pump strength conditions were characterized by a larger number of recruited nodes (Figure 5D, middle). As the Na^+^/K^+^ pump strength increased, the number of recruited nodes at the peaks in RSS reduced. Similarly, networks with lower pump strengths displayed stronger correlations between ETS-based FC and time-averaged rsFC (Figure 5D, right). As the Na^+^/K^+^ pump regulates [K^+^]_o_, lower pump strengths (e.g. 90% condition) allow for larger/slower [K^+^]_o_ fluctuations whereas higher pump strengths (e.g. 120%) lead to a tighter regulation of [K^+^]_o_ and smaller/faster fluctuations. Larger/slow [K^+^]_o_ fluctuations increase the window of opportunity for nodes in the network to synchronize or become active, thereby contributing to the RSS peaks. Alternatively, increasing the pump strength means that when [K^+^]_o_ begins to increase it is very quickly discharged and so the [K^+^]_o_ increase is short lived. This quick discharge reduces the window of opportunity for node recruitment and formation of RSS peaks.

**Figure 5.**
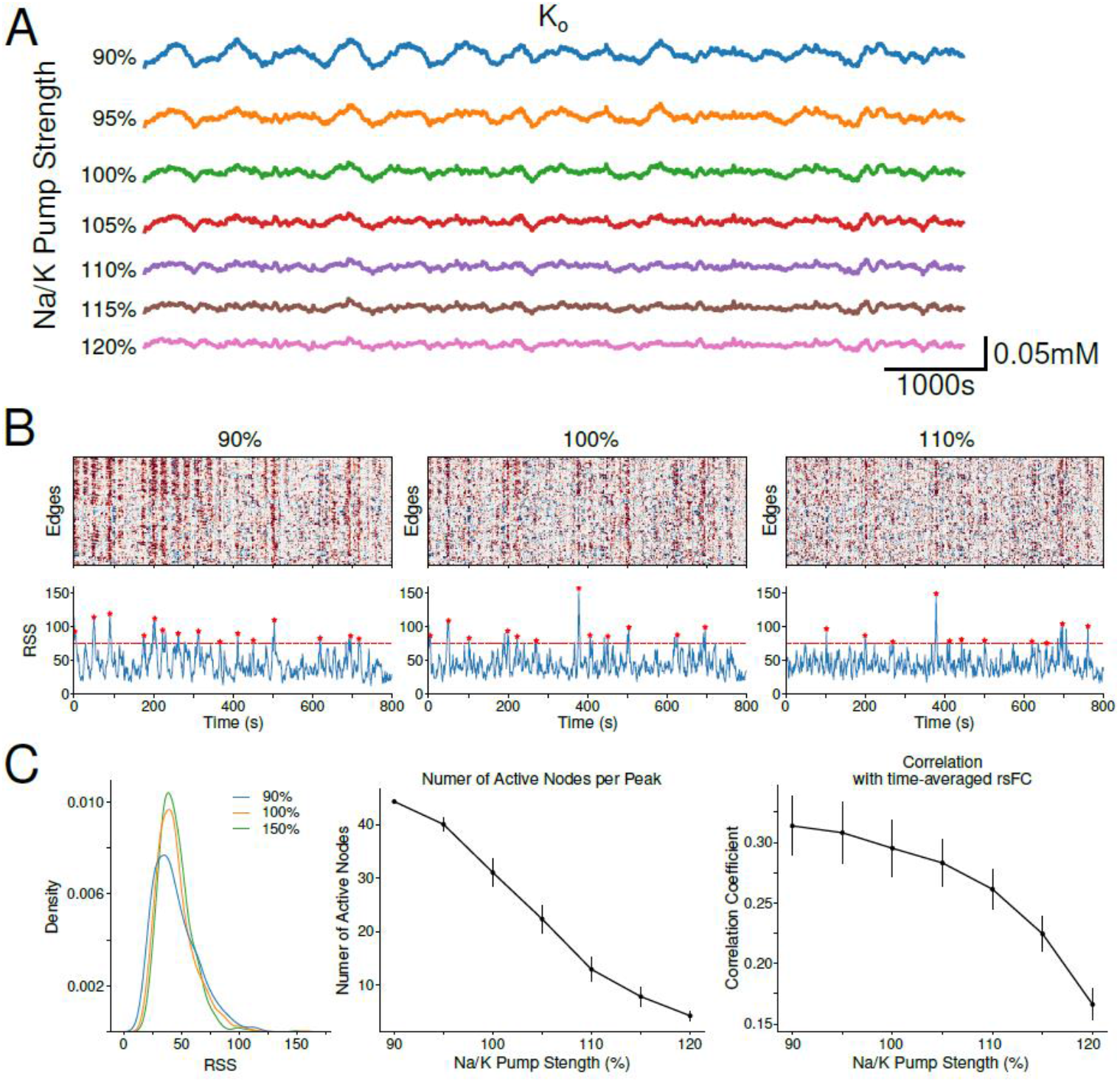
Decreasing Na^+^/K^+^ pump increases recruitment of nodes in tail RSS events. A, Mean [K^+^]_o_ of networks with varying amounts of Na^+^/K^+^ pump strengths. Percentages listed to the left indicate the percent strength of Na^+^/K^+^ pump with 100% being the default/baseline network condition. B, ETS computed for 3 network conditions 90% (left), 100% (middle), and 110% (right) Na^+^/K^+^ pump strengths. Corresponding RSS time series below each ETS. Red dashed line demarcates the threshold use to detect peaks and stars indicate detect peaks. C, Left, Distributions of RSS values in B. Middle, Average number of active nodes per detected RSS peak as a function of Na^+^/K^+^ pump strength. A node is considered active if their firing rate increases past its average firing rate during that RSS peak. Right, Correlation between time-averaged rsFC and ETS-based FC as a function of Na^+^/K^+^ pump strength. 95-percentile was used as a threshold for RSS peak detection.

### E-I balance determines the spatial recruitment during the events

In the previous section we explored how regulation of local ion concentration dynamics influences node recruitment and moment-to-moment co-fluctuations in activity underlying FC. We next asked how synaptic connectivity, either long-range or local, influenced these co-fluctuations and node recruitment. First, we explored the role of long-range connectivity in node recruitment. We varied the strength of excitatory feedforward connections between nodes in a network defined by human structural connectivity (Figure 1E). We found that increasing all feedforward long-range excitatory connections results in an increase in node recruitment at RSS peaks (Figure 6A, left). This increase in node recruitment was accompanied by increases in the amplitudes of RSS peaks (Figure 6A, right). Alternatively, we observed a negative correlation between long-range feedforward inhibition and node recruitment (Figure 6B, left). Increasing feedforward inhibition resulted in a slight decrease in the amplitude of RSS peaks (Figure 6B, right). These results suggest that long-range connectivity can influence co-fluctuations in activity by modulating recruitment of nodes and synchronization of the network as a whole. Next, we explored the role of local excitatory/inhibitory (E-I) balance on co-fluctuations. For these experiments, we kept feedforward connection strengths constant (i.e. 100% scaling) and only varied the excitatory/inhibitory synapses within each node. By doing this sweep in local E-I balance, we found that node recruitment at RSS peaks was highest in conditions described by low inhibition (Figure 6C, left). Interestingly, not all conditions where node recruitment was highest correlated with increases in the number of RSS tail events or peaks (Figure 6C, middle). We found that the highest number of tail events or RSS peaks occurred in conditions of high excitation and low inhibition (Figure 6C, middle). Similarly, the highest correlated activity across the network was seen in these same low inhibition / high excitation conditions (Figure 6C, right). These findings suggest that local E-I balance is crucial for correlated activity across the whole network and impacts moment-to-moment co-fluctuations underlying FC. Together, these results demonstrate the role of both local and long-range synaptic strengths on the properties of co-fluctuations in activity driving rsFC.

**Figure 6.**
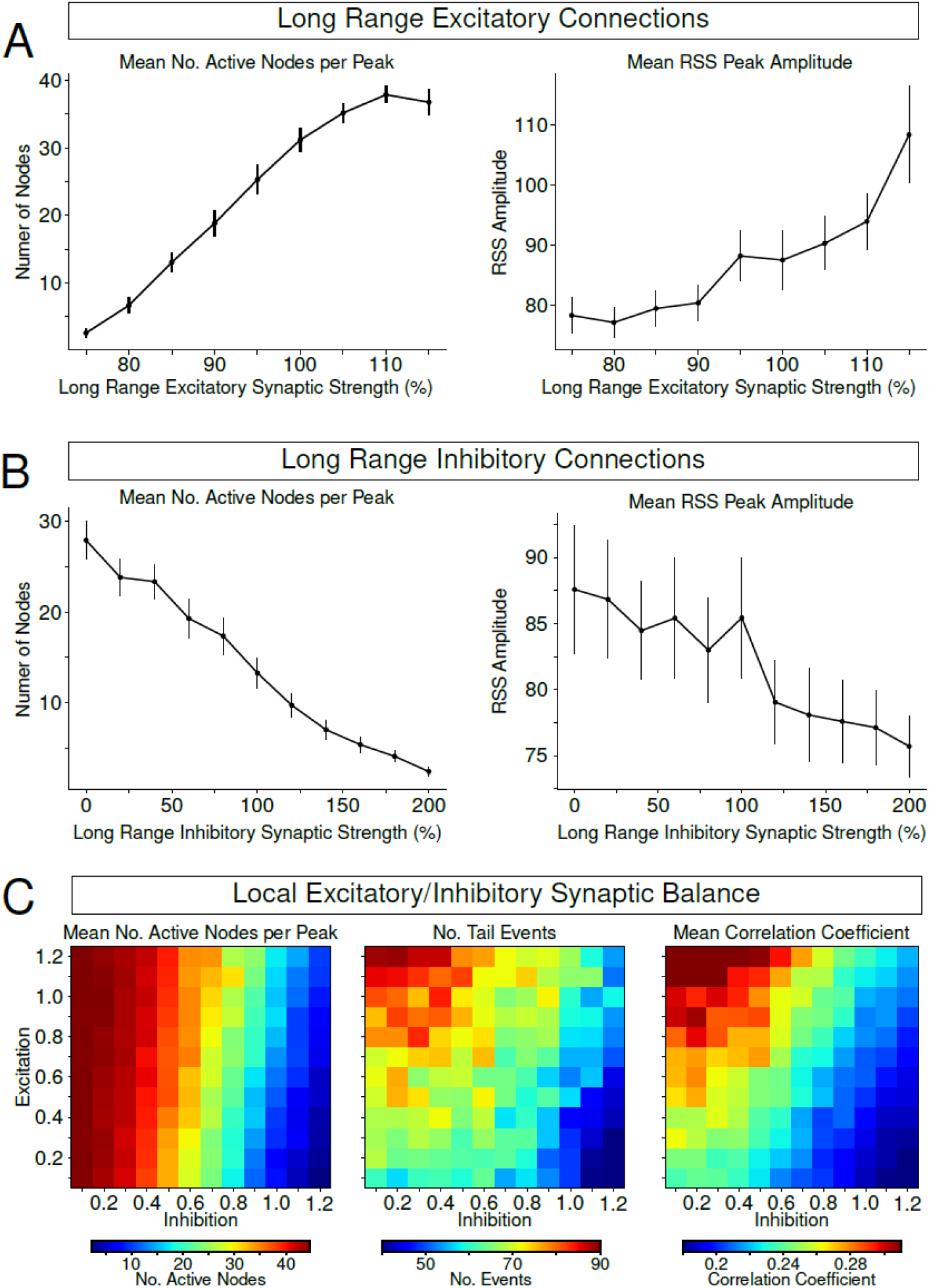
Feedforward excitation drives more nodes recruitment (and vice versa for inhibition). A/B, Left panel show the average number of active nodes per RSS peak as a function of the strength of long-range feedforward excitatory (A)/inhibitory (B) synapses between nodes. Right panel shows the average amplitude of the detected RSS peaks as a function of long-range feedforward excitatory (A)/inhibitory (B) synapses. C, Left heatmap shows the effect of changes in strength of local excitatory (y-axis) and local inhibitory (x-axis) synaptic strength on the average number of active nodes per RSS peak. Middle heatmap shows the resulting number of tail events identified from RSS distributions when varying local E-I balance. Right heatmap shows the mean functional connectivity computed across the entire network for different E-I balance conditions.

## Discussion

BOLD activity in the brain shows spontaneous infra-slow fluctuations (41). While there is ongoing discussion about the relative contribution of different physiological factors to the BOLD signal (7), there is ample evidence supporting a neural origin of infra-slow fluctuations (8-12, 42), and the derived static and time-varying functional connectivity (see (13) for a review). FC is typically measured using linear Pearson correlation coefficient, as the nonlinear contributions to coupling are practically negligible (43, 44). Notably, the FC is traditionally computed from the full time series, based on the assumption of stationarity and the need for sufficiently long recordings to obtain a reliable estimate of FC (45). Of course, alternative approaches have also been proposed, often centered on the idea of switching brain states characterized by distinct FC (13, 14, 46), although the inference of such states may be methodologically problematic (27, 47-49). In an extreme case, one could view brain dynamics as a progression of states defined by an instantaneous vector of brain activation across voxels, or regions, with the observed FC representing the time-averaged result of the observation of the covariance. From this perspective, stationary FC can be well approximated from a very sparse sampling of the original process, by considering only the timepoints with the largest amplitude (17, 18, 50). A similar observation has been formulated in the framework of edge-defined space (19, 51), which has drawn much attention and discussion (20, 22, 52-55), suggesting that such behavior may be expected for a broad range of processes while leaving the origin of these fluctuations an open question.

A complementary approach to studying experimental data with unknown ground truth is to model the possible neural ground truth and propose its potential origins, assuming the neural basis of the signal. Previous modelling work has already suggested possible mechanisms for the emergence of co-fluctuations and their patterns based on the network’s structural topology and the oscillatory nature of the underlying dynamics (either via mass models [e.g. (56)] or by using simple phase-oscillators (16, 57)). In this new work we offer a direct biophysical connection to the ionic-scale, linking the observed co-fluctuations to extracellular potassium levels and the associated Na^+^/K^+^ pump dynamics, which account for roughly half of brain energy consumption (58, 59) and thus contributing to the BOLD signal itself.

We use a detailed biophysical model of infra-slow fluctuations in resting-state (32) in the human connectome which reproduces FC seen in humans (60) and analyze co-fluctuations dynamics in the framework of (19). We observed brief periods of high co-fluctuations, which are driven by underlying ionic changes reflected in [K^+^]_o_, Na^+^/K^+^ pump current, firing rate and average synaptic input variables of the model. We report similar results for the macaque connectome. We show that potassium dynamics affects the occurrence of co-fluctuation events and fixing [K^+^]_o_ shifts inter-event intervals to a shorter range. Both increasing and decreasing neural patterns of firing and K^+^ levels were associated with different high co-fluctuations events. Clearance of [K^+^]_o_ is determined by the effectivity of the Na^+^/K^+^ pump, and we found that manipulating the pump strength influenced the co-fluctuations events not only in amplitude but also in spatial extent. This implies a changed network configuration in which more nodes have a chance to coordinate their activity.

Apart from ionic changes, both local and long-range E-I balance in connectivity influenced co-fluctuation events both in amplitude and node recruitment, though different E-I balance was required for maximizing amplitude or for node recruitment. We show that correlated activity across the network (i.e. functional connectivity) can be influenced by E-I balance as well, and was maximized for the low inhibitory - high excitatory condition. We conclude that low level ionic changes, which influence neural dynamics, propagate to BOLD signal and its co-fluctuations, and thus offer possible biophysical explanation underlying these events.

## Methods

### Ions dynamics and intrinsic excitability

Our computational model used in this study has been described in detail elsewhere (61-65). Briefly, excitatory (PYs) and inhibitory (INs) neurons were modeled as two-compartment neurons comprised of a dendritic and an axosomatic compartment. The temporal evolution of voltage for each compartment was described by the following equations:

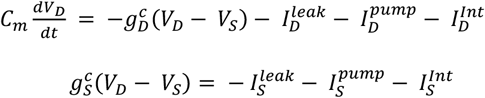

where *V*_*D,S*_ are the voltages of the dendritic and axosomatic compartments (respectively), 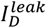 and 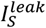 are the sum of the ionic leak currents, 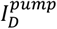 and 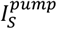 are the sum of the Na^+^ and K^+^ currents through the Na^+^/K^+^ pump, and 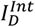 and 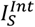 are the intrinsic currents for the dendritic and axosomatic compartments respectively. Each of these compartments contained conductance-based Hodgkin-Huxley type ionic currents. The axosomatic compartment contains the fast sodium current (*I*_*Na*_), the persistent sodium current (*I*_*NaP*_), delayed-rectifier potassium current (*I*_*Kv*_), and the sodium-activated potassium current (*I*_*KNa*_). The intrinsic ion currents in the dendritic compartment include the fast sodium (*I*_*Na*_), persistent sodium current (*I*_*NaP*_), slowly activating potassium current (*I*_*Km*_), high-threshold calcium current (*I*_*Ca*_), calcium-activated potassium current (*I*_*KCa*_), hyperpolarization-activated depolarizing mix cationic currents (*I*_*h*_), and leak currents (63-65). Na^+^/K^+^ pump and KCC2 cotransporter Cl^-^ extrusion were included in both neuron types. Additionally, ion concentration dynamics for extracellular and intracellular Na^+^ and K^+^ as well as intracellular Cl^-^ and Ca^2+^ were determined by the intrinsic currents, transporter-mediated currents, leak currents, extracellular and intracellular diffusion, and glial [K^+^]_o_ buffering as described in the following equations:

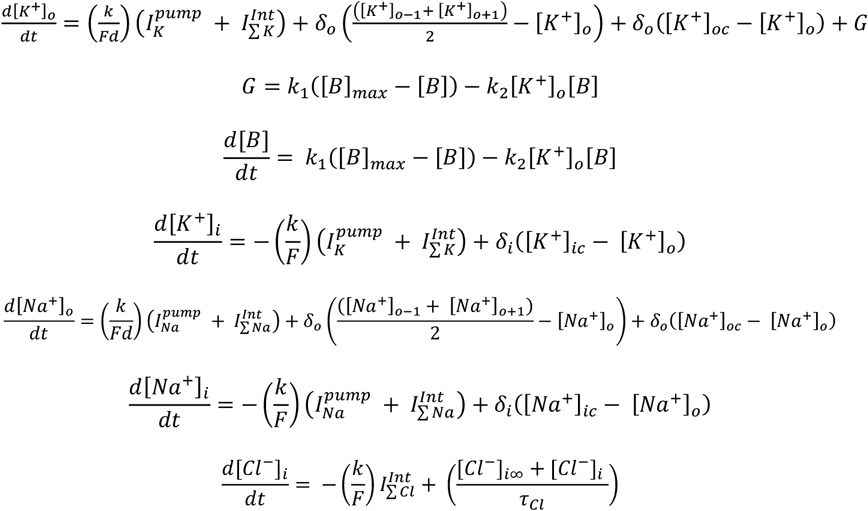

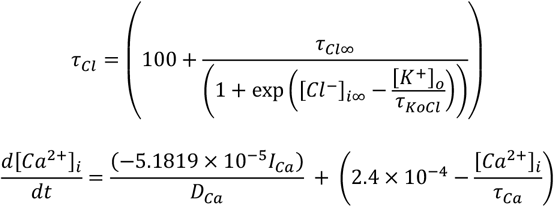

where *F* = 96489 C/mol, *d* = 0.15 is the ratio of the extracellular compartment volume to surface area, the conversion factor *k* = 10, *δ*_*o*_ is the scaled diffusion coefficient (*δ*_*o*_ = *D*/Δ*x*) where *D* = 6×10^-6^ *cm*^2^/*s* is the diffusion constant and Δ*x* = 100 *μm* is distance, [*K*^+^]_*oc*_ and [*Na*^+^]_*oc*_ are the K^+^ and Na^+^ concentrations in the adjacent compartments, and [*K*^+^]_*o*−1_, [*K*^+^]_*o*+1_, [*Na*^+^]_*o*−1_, and [*Na*^+^]_*o*+1_ are the concentrations of K^+^ and Na^+^ in neighboring cells respectively. Astrocytic glial buffering of extracellular K^+^ (*G*) was modeled as a free buffer ([*B*]) with total buffer ([*B*]_*max*_) = 500mM. The [*B*] K^+^ binding and unbinding rates (*k*_1_ and *k*_2_ respectively) were given by

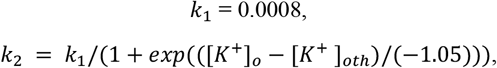

where [*K*^+^]_*oth*_ = 15mM is the half activation concentration of [*K*^+^]_*o*_. [*Cl*^−^]_*i*∞_ = 5mM, *τ*_*Cl*∞_ = 2×10^4^, and *τ*_*KoCl*_ = 0.08s. *τ*_*Ca*_ and *D*_*Ca*_were set to 300ms and 0.85 respectively. Extracellular K^+^ was also allowed to diffuse between the two compartments as well as between neighboring cells of the same type (ie. diffusion between PY-PYs and IN-INs). Some slow time constants can be found in our equations for glial K^+^ buffering and Cl^-^ transport. However, these slow rate constants are faster than the observed infra-slow time scale of the neural dynamics arising in our network.

### Synaptic properties and local network connectivity

Each local cluster or individual brain region in our model was comprised of 50 PY and 10 IN neurons. Each PY neuron made local excitatory connections onto 10 other PY neurons and received 10 excitatory connections from other PY neurons. PY neurons also formed excitatory connections onto inhibitory IN neurons. Each PY projected onto one IN and each IN formed inhibitory connections onto 5 PY neurons. Excitatory connections were mediated by AMPA and NMDA conductances (11 nS and 1 nS, respectively), and inhibitory connections were mediated by GABA_A_ conductances (11 nS) such as those described previously (63-65). Excitatory connections from PY neurons onto IN neurons were mediated by AMPA and NMDA conductances (3.5 nS and 0.35 nS, respectively). To model *in vivo* conditions, all neurons of both types received additional afferent excitatory input as a random Poisson process.

### Macaque connectivity

We implemented the structural connectivity of 58 macaque brain regions. Connection strengths between brain regions were extracted from the CoCoMac database (http://cocomac.g-node.org). Functional connectivity was computed as the correlation coefficients between mean Na^+^/K^+^ pump currents or computed BOLD signals from individual clusters. Significance values were Bonferroni corrected to correct for multiple comparisons.

### Human connectivity

Acquisition of MRI data and construction of structural connectivity was identical to the methods described in (36). To summarize, the data provided here are based on MRI scans of 90 healthy control individuals. The construction of structural connectivity matrices was based on a connectome generated by probabilistic tractography on diffusion MRI data. We used ROIs from the widely used AAL atlas (66). The connectivity between two ROIs was based on the number of streamlines in the tractogram beginning in one ROI and terminating in the other ROI. Global coupling and the produced baseline dynamics were identical to the one used in (60).

## Acknowledgements

O.C.G. is supported by NIA award K99AG086609. PS and JH were supported by ERDF-Project Brain dynamics, No. CZ.02.01.01/ 00/22_008/0004643 and by the Czech Science Foundation project No. 21-32608S. GPK and MB were partially supported by NSF award 2209874. MB was supported by NIH award 1R01MH125557, 1RF1NS132913.

## Supplementary Figure

**Figure S1.**
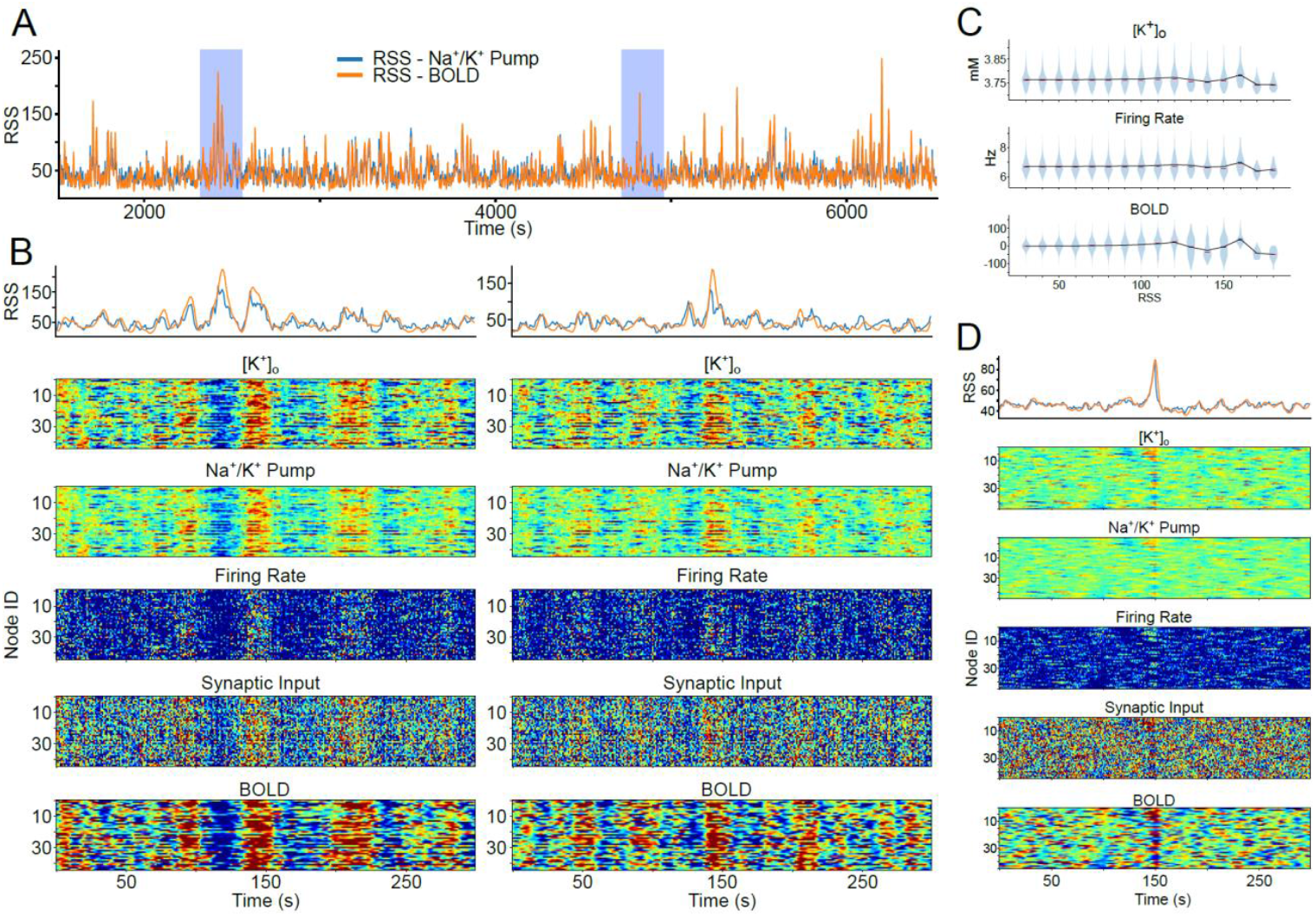
A.Trace of edge-based time series estimated for entire human connectome network for 1000sec simulation. B. Zoom-in of two periods when the RSS had values above 95th percentile. From top, the plots show the RSS of the whole network, extracellular K^+^, Na^+^/K^+^ pump, firing rate, average synaptic input and BOLD across individual regions for the selected period. C. Violin plot show the distribution of [K^+^]_o_ (top), firing rate (middle) and BOLD(bottom) for bins of RSS value from the entire simulation period. D. RSS event (defined as RSS > 95th percentile) triggered average of various neural measures across different regions within 150 sec window of the event.

